# Symbiodiniaceae conduct under natural bleaching stress during advanced gametogenesis stages of the mesophotic coral *Alveopora allingi*

**DOI:** 10.1101/2021.03.25.437087

**Authors:** Gal Eyal, Lee Eyal-Shaham, Yossi Loya

## Abstract

The mesophotic coral *Alveopora allingi* from the northern Gulf of Eilat/Aqaba, Red Sea, is affected by year-round partial coral-bleaching events. During these events, the migration of Symbiodiniaceae takes place from the coral-host mesoglea to the developed oocytes in bleached parts of colonies of *A. allingi* but not in the non-bleached parts. Additionally, these oocytes are abnormal, missing part of the structural material of the peripheral areas and are also significantly larger in the bleached areas of the colonies. Hence, we suggest a parasitic behavior of the symbionts or a commensalism relationship which enhance symbionts’ needs during bleaching periods and may boost the gametogenesis development in these corals. We propose that evolutionarily, this behavior may greatly contribute to the symbiont community survival throughout the bleaching period, and it can also be beneficial for the host’s persistence and adaptation to bleaching through the acquisition of a specific symbiont community following the bleaching event.

## Introduction

Coral reefs have experienced increasing stress over the last four decades due to local anthropogenic perturbations and global climate change, resulting in severe damage to coral populations and their reproduction (Loya 2004,2007; Carpenter et al. 2008; Baird et al. 2009; Harrison 2011; Hughes et al. 2017). The symbiotic relationship between corals and their algal endosymbionts (Symbiodiniaceae) is a key factor in the evolutionary success of hermatypic corals (Wooldridge 2010). This close association between primary producer and consumer enables the tight nutrient recycling that is thought to explain the high productivity of coral reefs (Hoegh-Guldberg 1999; Wooldridge 2013; Muller-Parker et al. 2015). Consequently, environmental and physiological conditions that result in changes in the relationship between animal host and Symbiodiniaceae may have profound ecological and physiological effects (Szmant and Gassman 1990). The loss of Symbiodiniaceae - a phenomenon described as coral bleaching - can cause acute damage to the colony (Glynn 1993; Brown 1997; Loya et al. 2001; Suggett and Smith 2020). Coral bleaching has been observed in response to a diverse range of stressors and is associated with both anthropogenic and natural disturbances (Fitt and Warner 1995; Glynn 1996; Kushmaro et al. 1996; Hoegh-Guldberg 1999; Rowan 2004; Hughes et al. 2017; Hughes et al. 2018). Severe and prolonged bleaching can cause partial to total colony death, resulting in diminished reef growth, transformation of reef-building communities to degraded alternate states, increased bioerosion and, ultimately, the disappearance of reef structures (Glynn 1996). The future state of shallow reefs is glooming and the framework building of coral communities is expected to shift over to low-relief and less complex entities (Knowlton 2001; Pandolfi et al. 2003; Hughes et al. 2007).

While shallow reefs have recently been intensively affected by mass coral-bleaching (Hughes et al. 2018), very few mesophotic coral ecosystems (MCEs; 30-150 m depth) were reported to have undergone bleaching during these events (Baker et al. 2016; Frade et al. 2018). One explanation could be that of the ‘deep reef refuge’ hypothesis (DRRH), which suggests that deep ecosystems are more protected from the disturbances that affect shallow reefs, and consequently could provide a viable reproductive source for shallow-reef areas following disturbance (Bongaerts et al. 2010). MCEs too, however, are not immune to bleaching (Frade et al. 2018) or to other disturbances (Rocha et al. 2018; Pinheiro et al. 2019), which in many cases have been overlooked due to technical difficulties. One of the most important questions facing scientists, policy-makers, and the general public is that of why there has been an apparent increase in the incidence of coral bleaching in the last four decades (Goreau and Hayes 1994; Hoegh-Guldberg 1999; Hughes et al. 2017) and what we can do about it?

Nevertheless, although highly impacted by anthropogenic stressors, the shallow corals of the Gulf of Eilat/Aqaba (GoE/A) (northern Red Sea) are considered heat-tolerant and resilient to thermal stress (Fine et al. 2013; Krueger et al. 2017; Osman et al. 2018). They have been described as comprising ‘super-corals’ that do not experience natural bleaching (Grottoli et al. 2017). Nonetheless, *Stylophora pistillata,* the most abundant depth-generalist coral in the coral reefs of Eilat, suffers from periodic non-fatal bleaching in the summer months in the upper MCEs (Nir et al. 2014), and recent observations have revealed multiple-species bleaching-events on these deeper ecosystems (Eyal et al. 2019).

Symbiotic relationship between the coral host and its Symbiodiniaceae community are complex and essential for the development of reef corals (Trench 1971; LaJeunesse 2020). Nevertheless, these relationships are not necessarily always beneficial for both the host and the symbionts. Parasitism in cnidarian symbiosis under stress conditions was suggested in the past but supported only by indirect evidences (Baker et al. 2018; Peng et al. 2020). Here we use observations from coral reproductive study to present another potential hypothesis of parasitism of the symbionts or/and commensalism relationship between the host and its symbionts.

Sexual reproduction is the most critical part of any living taxa (Kondrashov 1988). Although energetically costly, the benefit of increasing fitness and genetic diversity is priceless. Evolution processes through sexual reproduction and gene-shuffling is the primary adaptive capability to environmental changes and enabling extension to and occupation of new ecological niches (Rundle et al. 2006). In shallow corals, diverse strategies of sexuality were discovered during recent years (Baird et al. 2009; Harrison 2011; Eyal-Shaham et al. 2019; Eyal-Shaham et al. 2020) and references within) but our knowledge of the reproduction of MCE corals remains limited (Shlesinger and Loya 2019).

The purpose of this research was to detect possible natural bleaching effects on the development of gametogenesis in mesophotic corals. Hence, we examine the reproductive ecology of the mesophotic coral, *A. allingi,* under bleaching stress and describe a novel host-symbiont behavior interaction under bleaching conditions.

## Methods

### Study area, sampling procedures, and histological processes

Coral pairs from the same genets of non-bleached (pigmented) and visually bleached *A. allingi* were sampled in front of the Interuniversity Institute for Marine Sciences in Eilat, Israel (IUI) (29°30′N, Θ34°55′E), at a depth of 60 m, during the end of their 2011 reproductive season (October 2011). This is a well-developed reef with a wide variety of coral species, and *A. allingi* is one of the most abundant species in the area (Eyal-Shaham et al. 2016). As part of a larger project described in Eyal-Shaham et al. (2016), once a month, during the full moon, nubbins (4–5 cm in length) were randomly sampled from 4–6 healthy colonies of *A. allingi* using SCUBA technical diving and Closed-Circuit Rebreathers (CCR). Only in October 2021 we found partially bleached colonies (two different genet pairs of pigmented and visually bleached nubbins). The sampled colonies were >20 m distant from one another. The collected nubbins were placed in a sealed nylon bag filled with seawater and immediately after collection were fixed in a 4% formaldehyde solution in seawater for 24–48 h. The nubbins were then rinsed in tap water for 15 minutes and transferred to 70% ethyl alcohol. The decalcification process was carried out following the protocol of Eyal-Shaham et al. (2016). A piece of tissue was taken from each decalcified nubbin and 6 μm thick latitudinal histological serial sections were prepared and dyed with Mayer’s hematoxylin and Putt’s eosin (H&E stain) to highlight the reproduction structures.

### Histological and statistical analyses

The histological analysis was performed using a Nikon Eclipse 90i microscope and NIS Elements D 3.2 software (Nikon Instruments Inc.). A detailed study of gonad structure, size, and development was conducted by analyzing the histological cross-sections [see details in Eyal-Shaham et al. (2016)]. Detailed identification of the oocytes (Oc), spermaries (Sp), nucleus and nucleoli (N), and Symbiodiniaceae (Z) were conducted. The examination included size measurements of the oocytes (longest axes), performed only when the nucleoli were present, and identification of the developmental stages of gametocytes within the polyps. Mean oocyte size of each developmental stage was calculated from all oocytes presented in the samples from the bleached and the healthy groups.

Differences in oocyte sizes within partly bleached colonies were calculated in R language and environment (R Development Core Team 2014) using 1,000 bootstrap samples of the values to indicate the difference between the median oocyte size of the bleached part and that of the control part.

## Results and Discussion

During coral bleaching events that occur at 50-60 m depth, the depth-specialist *A. allingi* experiences severe visual bleaching in parts of the branches of otherwise healthy colonies (Fig. 1a, b). Unlike the periodic summer bleaching of *S. pistillata,* also described from the same reef (Nir et al. 2014), and in which the colonies recover after the summer ends, *A. allingi* presents a gradual degradation of the bleached tissue, with mortality of the entire bleached part of the colonies occurring within a few months.

**Figure 1:**
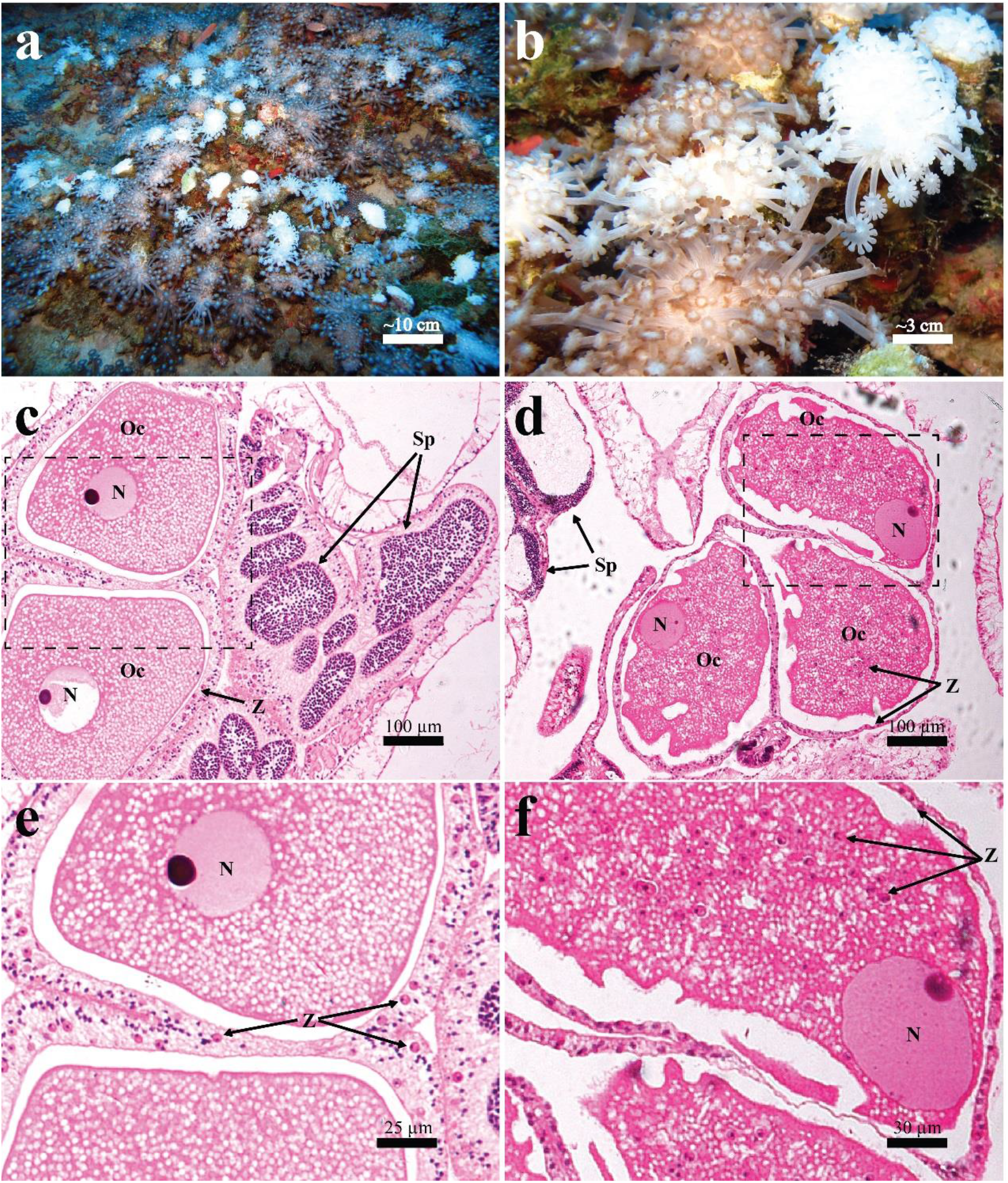
Partially-bleached *Alveopora allingi*, with micrographs of gonadal development. (a) general colony view at 60 m depth; (b) partial-bleaching within a single genet; (c) male and female gonads from the non-bleached (pigmented) part of the colony; (d) male and female gonads from the visually bleached part of the colony; (e) magnification of the marked area in section c, showing stage 4 healthy mature oocytes; and (f) magnification of the marked area in section (d), showing several stage 4 severely damaged oocytes with many Symbiodiniaceae in the oocytes’ cytoplasm. Scale bars as indicated above the scale-line. Oc: Oocyte, Sp: Spermaries, N: nucleus, Z: Symbiodiniaceae.

Our analysis of the histological sections of partly-bleached colonies revealed a phenomenon that to our knowledge has never previously been described: Symbiodiniaceae migration to the oocyte cytoplasm [i.e., in the visually bleached part of the colonies, Symbiodiniaceae were observed in the oocytes’ cytoplasm, which presented an irregular shape and imperfect cytoplasm with missing parts of the structural material of the peripheral areas, while the healthy (non-bleached) portions of the same colonies presented normally-developed oocytes (Fig. 1c-f)]. Additionally, the bleached coral’s oocytes were found to be significantly larger than those of the non-bleached parts (Fig. 2; Mann-Whitney Test, T=544, P<0.001), which suggests a faster reproduction development in the impacted parts of these colonies. Although Symbiodiniaceae can be directly transmitted from the parent colony (i.e. vertical transmission) in some species (Lesser et al. 2013; Muller-Parker et al. 2015), the observed change in shape (i.e., larger sizes and imperfect cytoplasm with missing parts of the structural material of the peripheral areas of the oocytes only), suggests that this is not the case in *A. allingi.* Moreover, Symbiodiniaceae were not observed in any mature oocytes of other, healthy, colonies.

**Figure 2:**
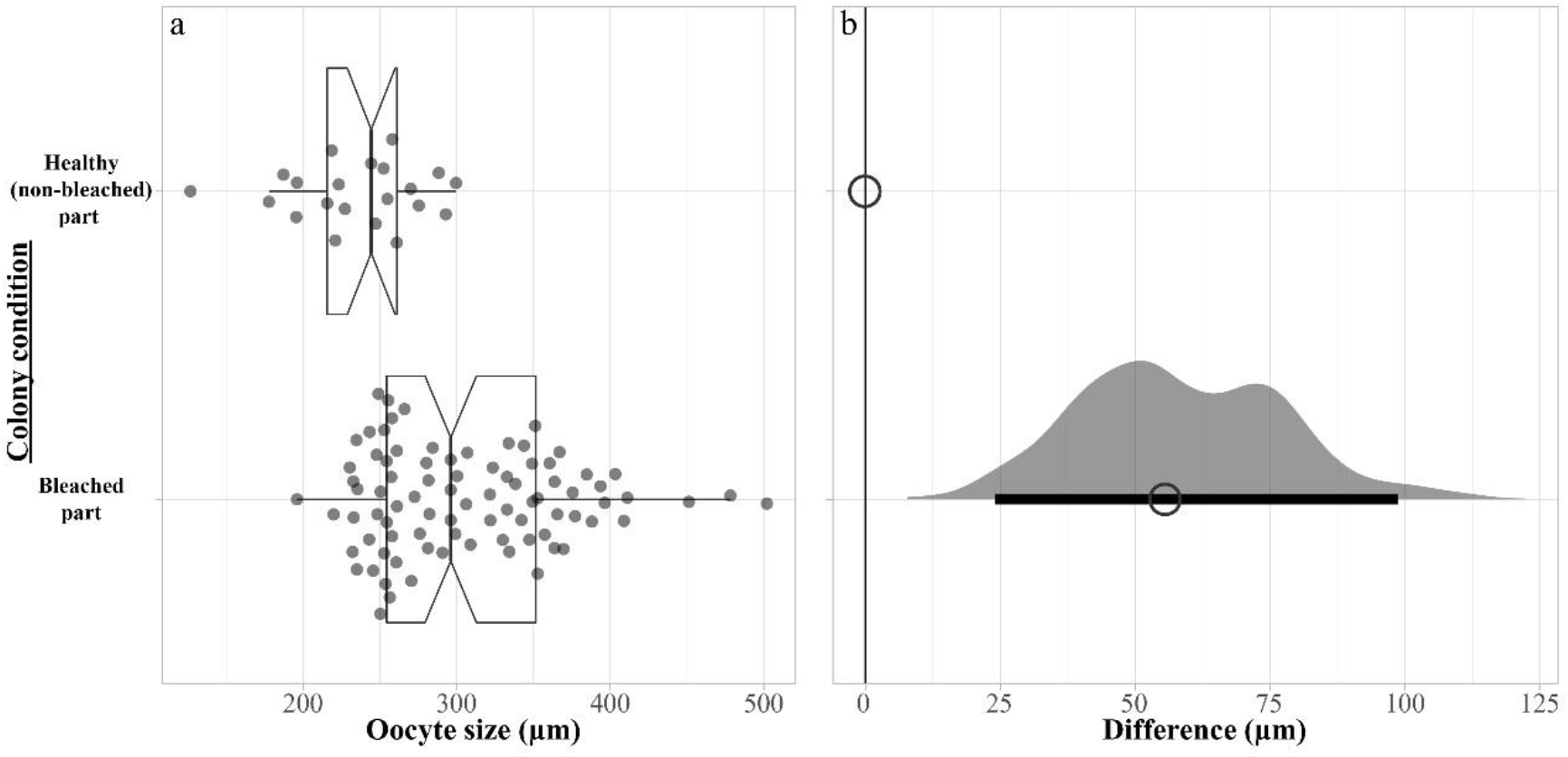
Oocyte sizes of the mesophotic coral *Alveopora allingi* during a partial colony natural bleaching event. (a) Significantly larger stage 4 oocytes in the bleached parts of partly bleached colonies (t-test, T=544, P<0.001). Centerlines of the boxes represent the medians; box limits indicate the 25th and 75th percentiles; whiskers extend 1.5 times the interquartile range from the 25th and 75th percentiles, notches represent the 95% confidence interval for each median, all data represented by dots. Two different genet pairs of bleached (n=84 oocytes) and non-bleached (n=21 oocytes) parts were analyzed. (b) The difference in μm between median values of the bleached parts compared to non-bleached parts. The difference in condition is calculated from 1,000 bootstrap samples of the values, indicating the difference between the median oocyte size of the bleached parts and that of the healthy parts. The horizontal black line indicates the 95% confidence interval.

We hypothesize that such migration of Symbiodiniaceae into the coral oocyte cytoplasm under severe bleaching conditions potentially enhances the survivorship of the symbiont community, suggesting the existence of a possible mechanism of exploitation of cytoplasmic materials by certain symbionts. An alternative hypothesis is that the partially bleached hosts are in a process of absorbing oocyte lipids as an emergency energetic source. Cell membranes have been disturbed, including the symbiosome surrounding the host cells lining the exterior of the oocyte. This enables symbionts to escape the host cell and invade the oocytes.

The Symbiodiniaceae community in corals is generally dominated by one or two main symbiont types (van Oppen et al. 2005; Ziegler et al. 2017) but can also contain a large number of rarer members of the family, which play important roles in the resilience of the holobiont (Ziegler et al. 2018). Network theoretic modeling predicts that elevated symbiont diversity can increase community stability in response to environmental changes (Fabina et al. 2013). Moreover, the functional role of Symbiodiniaceae in the holobiont is not defined by a single species but rather by the assemblages of symbionts (Ziegler et al. 2018). The rare symbionts, however, associate in several manners with the coral host (Knowlton and Rohwer 2003; Kirk et al. 2013; Baker et al. 2018), which may include non-mutualistic species. Additionally, some Symbiodiniaceae may transform their mutualistic relationship into a parasitic state during a stressed period, thus parasitizing their hosts for nutrition (Baker et al. 2018). The symbiont benefits are clearly not equal to those of the host, host and this partnership could be lost under times of stress to either the host or the symbionts. In addition to this novel behavior- the phenomenon of symbionts taking advantage of the stressed hosts, might also hold a key to some urgent questions regarding the coral-algae symbiosis breakdown and contribute to our understanding of the mechanisms involved in the bleaching phenomenon. The discovery that Symbiodiniaceae migrate into the oocytes of bleached parts of partially-bleached MCE corals, but not into the healthy (non-bleached) parts, reveals the existence of opportunistic demeanor of the symbionts on their coral host’s gametes, which may result in lowered functionality of the affected host parts. From an evolutionary point of view, this behavior is highly beneficial to the symbionts’ survival throughout the bleaching period, since expelled symbionts are not expected to survive for long in the environment (Hill and Ralph 2007); but could also be beneficial for the host’s persistence and adaptation to bleaching. Selection processes towards bleaching-resistant capabilities at the gamete level of heathy oocytes will result, in the long run, in adapted populations of this coral.

## Acknowledgements

This is a preprint of an article published in *Coral Reefs.* The final authenticated version is available online at: https://doi.org/10.1007/s00338-021-02082-1

We thank the Interuniversity Institute for Marine Sciences in Eilat for the logistical support, and Barbara Colorni for help with the histological work. The comments of two anonymous reviewers greatly improved the manuscript. This study was supported by the Israel Science Foundation (ISF) No. 1191/16 to YL and by the European Union’s Horizon 2020 research and innovation program under the Marie Skłodowska-Curie post-doctoral grant agreement No. 796025 to GE.

## Compliance with ethical standards

All samples were collected and treated according to the Israeli Nature and Parks Authority permit no. 2011/38249.

